# Effects of phage variation on Shiga toxin 2 (Stx2) production and the virulence of Stx-producing *Escherichia coli*

**DOI:** 10.1101/2024.03.06.583675

**Authors:** Keiji Nakamura, Haruyuki Nakayama-Imaohji, Munyeshyaka Emmanuel, Itsuki Taniguchi, Yasuhiro Gotoh, Junko Isobe, Keiko Kimata, Yukiko Igawa, Tomoko Kitahashi, Yohei Takahashi, Ryohei Nomoto, Kaori Iwabuchi, Yo Morimoto, Sunao Iyoda, Tomomi Kuwahara, Tetsuya Hayashi

## Abstract

Shiga toxin (Stx)-producing *Escherichia coli* (STEC) causes serious gastrointestinal illness, including hemorrhagic colitis and hemolytic uremic syndrome. Although all known Stxs (Stx1 and Stx2) are encoded by bacteriophages (Stx phages), the production of Stx2 is known to be a major risk factor for severe STEC infections. The production of Stx2, but not Stx1, is tightly coupled with the induction of Stx phages, and Stx2 production levels vary between STEC strains, even within the same serotype. Here, we analyzed the genomic diversity of all Stx phages in 71 strains representing the entire O145:H28 lineage, one of the major STECs. Our analysis revealed the highly dynamic nature of the Stx phages in O145:H28, including the independent acquisition of similar Stx phages by different sublineages and the frequent changes in Stx phages in the same sublineages due to the gain and loss of Stx phages. Analyses of Stx2 production levels in O145:H28 strains and K-12 lysogens of Stx2 phages of specific groups and types, which were defined by their early region sequences and CI repressors, respectively, revealed that short-tailed Stx2a phages (S-Stx2a phages) confer significantly greater Stx2 production to host strains than long-tailed Stx2a phages (L-Stx2a phages). However, L-Stx2a phages that encode a specific type of CI repressor promoted Stx2 production, comparable to the level of production among S-Stx2a phages, as well as promoted virulence to host strains, exceeding the level among other L-Stx2a phages. We also showed a clear link between the phage induction efficiency, which was primarily determined by the early region of each phage, and the level of Stx2 production by host strains. These results provide important insights into the diversification and dynamism of Stx phages and the relationship between the variations in Stx2 phages and the amount of Stx2 production by their host strains.

**Author summary:** Shiga toxin (Stx)-producing *Escherichia coli* (STEC) is an important human intestinal pathogen that causes severe illnesses. These bacteria produce Stx1, Stx2 or both toxins, but the production of Stx2 is an important measure of the virulence of STEC strains. While both types of Stx are encoded by bacteriophages (Stx phages), Stx2 production is tightly coupled with phage induction, and variations in Stx2 phages have been associated with variations in Stx2 production levels by their host O157:H7 STEC strains. However, in non-O157 STEC strains, the variation in Stx phages and its association with host strain production of Stx2 have not yet been fully analyzed. This systematic study of Stx phages in O145:H28 STEC reveals not only the marked genomic diversity and dynamism of Stx phages in this STEC lineage but also that short-tailed Stx2 phages and a specific group of long-tailed Stx2 phages induce high levels of Stx2 production by host strains, and this increased production is linked to the efficient induction of phages.

## Introduction

Shiga toxins (Stxs) are the key virulence factors of Stx-producing *Escherichia coli* (STEC), which causes diarrhea and hemorrhagic colitis with life-threatening complications, such as hemolytic uremic syndrome. Stxs are classified as Stx1 or Stx2, each of which include several subtypes (Stx1a, Stx1c-Stx1e; Stx2a-Stx2l) [1,2]. While STEC strains produce one or more Stx subtypes [3,4], epidemiological studies suggest that Stx2-producing strains cause more severe STEC infections than strains producing only Stx1 [5,6].

The *stx* genes are encoded by bacteriophages (Stx phages), and STEC strains acquire these genes via the lysogenization of Stx phages. Although the integration sites and genome sequences of Stx phages are highly variable even within the same serotype [7,8], Stx phages are morphologically divided into two types based on their tail structures, which are defined by late genes: lambda-like long-tailed phages (L-phages) and short-tailed phages (S-phages), similar to the Stx2a phages of O157:H7, such as Sp5 and 933W [9–15]. A total of 99% of 279 publicly available Stx phages can be classified into either type based on genetic content [16]. Both types of Stx phages encode several lambda-like regulator genes that modulate early and late gene expression, such as the *cI* and *q* genes [9,17–19]. While *cI* genes exhibit notable sequence diversity between Stx phages [20], how the variation in *cI* genes is associated with tail type is not known.

The *stx* genes are located downstream of the late gene promoter [9,19,21,22]. The expression of *stx1* is primarily under the control of the iron-regulated authentic promoter [23], although prophage induction-dependent production of Stx1 has been described in some O26:H11 strains [24,25]. In contrast, the expression of *stx2* depends on the late promoter [26,27], and Stx2 production is tightly coupled with phage induction; thus, variations in Stx2 phage genomes can affect the amount of Stx2 production by each strain, as has been shown in several STEC lineages [28–31]. For example, the variation in Stx2 production levels in O157:H7 STEC strains was associated with the subtypes of Stx2a phages (all are S-phages), as defined by their early regions [28]. In particular, in O157:H7 clade 8, a highly pathogenic lineage of O157:H7 STEC, the γ subtype of the Stx2a phage confers increased Stx2 production and pathogenicity to host strains than do the other clade 8 strains that carry the δ subtype of the Stx2a phage [30].

O145:H28 is one of the major non-O157 STEC serotypes [32]. We previously analyzed the whole-genome sequences (WGSs) of 239 O145:H28 strains, including a systematic analysis of the prophages in seven finished genomes, and revealed notable variations in the sequences and integration sites of Stx phages among O145:H28 strains [33]. In that study, we found that although the distribution of *stx1a* genes was biased toward specific clades, the *stx2a* genes were widely but variably distributed throughout the entire O145:H28 lineage. However, the precise variation in Stx phages and its impact on Stx production by host strains of this lineage have not been elucidated. In the present study, to address these clinically important issues, we performed a systematic analysis of Stx phages of STEC O145:H28 and analyzed the diversity and dynamism of Stx phages in O145:H28 and the association of Stx2 phage types with Stx2 production by host strains. Furthermore, using two strains, one carrying duplicated Stx2a phages and the other carrying two different types of Stx2a phages and their mutants lacking either of the two Stx2a phages, we analyzed the effect of the Stx2a phage copy number and the difference in phage type on Stx2 production and the virulence of host strains in isogenic backgrounds.

## Results

### Strain set and Stx phages

For detailed analyses of Stx phages in O145:H28 strains, we selected 64 strains from the 239 strains analyzed in our previous study [33], 59 of which were sequenced in our laboratory. This set included eight finished and 56 draft genomes and covered seven of the eight clades in ST32 (clades A-H) and the ST137/ST6130 lineage in O145:H28 (ST32 clade D strains were not available in our laboratory). Among the 56 strains for which only draft genomes were available, two were subjected to Nanopore long-read sequencing to obtain finished sequences by hybrid assembly. Seven genome-finished strains recently deposited in the NCBI database [34,35] were also included in the dataset (Table S1). Thus, the final set included 71 strains (Fig. 1 and Table 1), of which 48 were isolated in Japan and the remainder were isolated in the USA, Belgium, or Canada. Most strains were human isolates, but five bovine and four environmental/food isolates were included. There were five *stx* genotypes, and four strains carried two copies of the *stx2a* gene (Table 1).

**Fig. 1.**
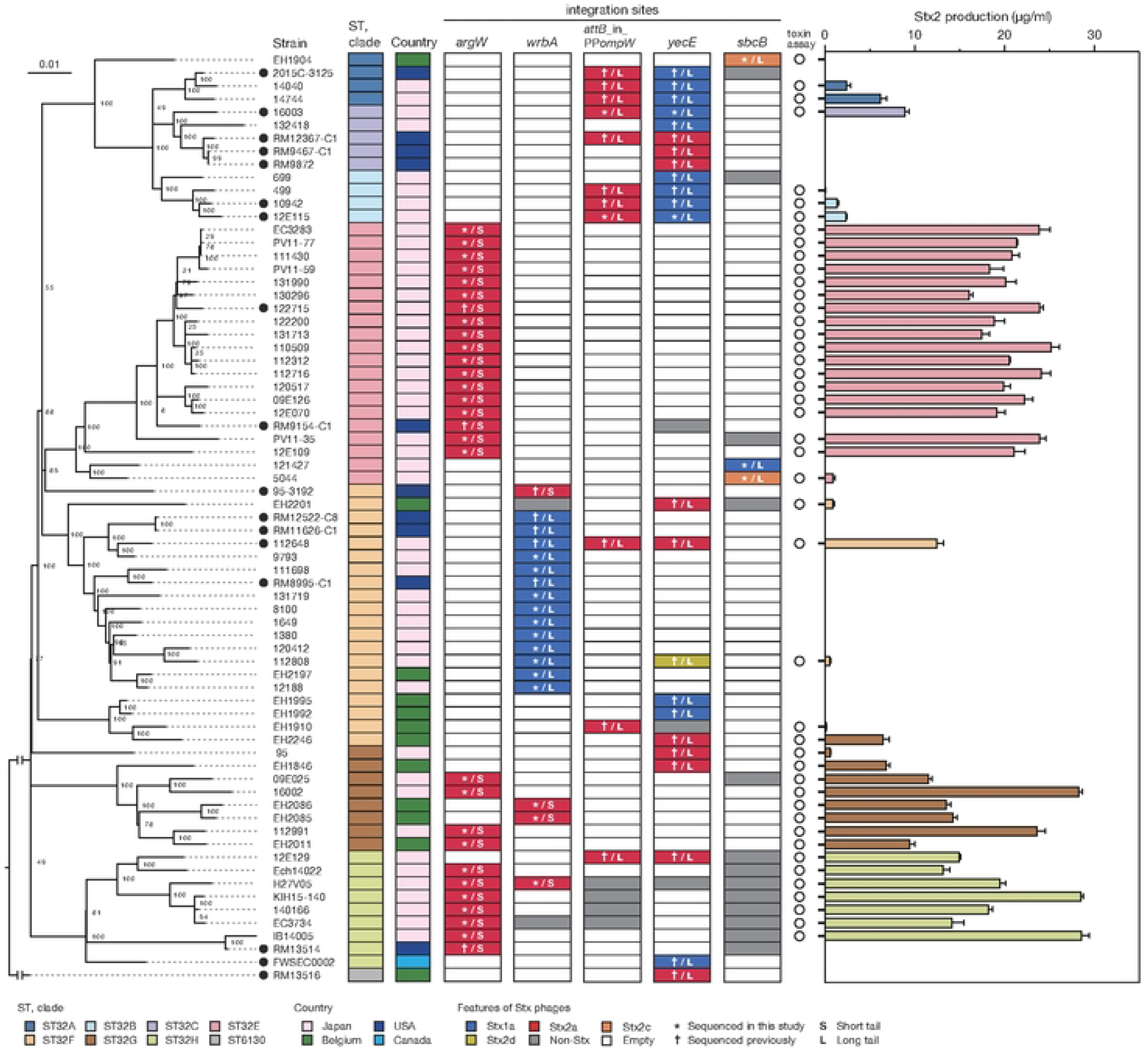
Variation in the integration site of Stx phages and the Stx2 production level in STEC O145:H28 strains. The phylogenetic tree of 71 O145:H28 strains is shown in the left panel. The tree was constructed based on the recombination-free SNPs (3,347 sites) that were identified on the conserved chromosome backbone (3,851,013 bp) by RAxML using the GTR gamma substitution model. The reliabilities of the tree’s internal branches were assessed by bootstrapping with 1,000 pseudoreplicates. The bar in the upper-left corner indicates the mean number of nucleotide substitutions per site. Genome-finished strains are indicated by filled circles. Along with the tree, the geographic and ST/clade information of strains, the presence or absence of prophages at five loci, and the features of prophages are shown. In the right panel, the levels of MMC-induced Stx2 production by each strain are shown as the mean values with standard errors of biological triplicates. Note that the Stx2 production levels of eight *stx2*-positive strains whose genome sequences were obtained from NCBI were not determined.

**Table 1.**
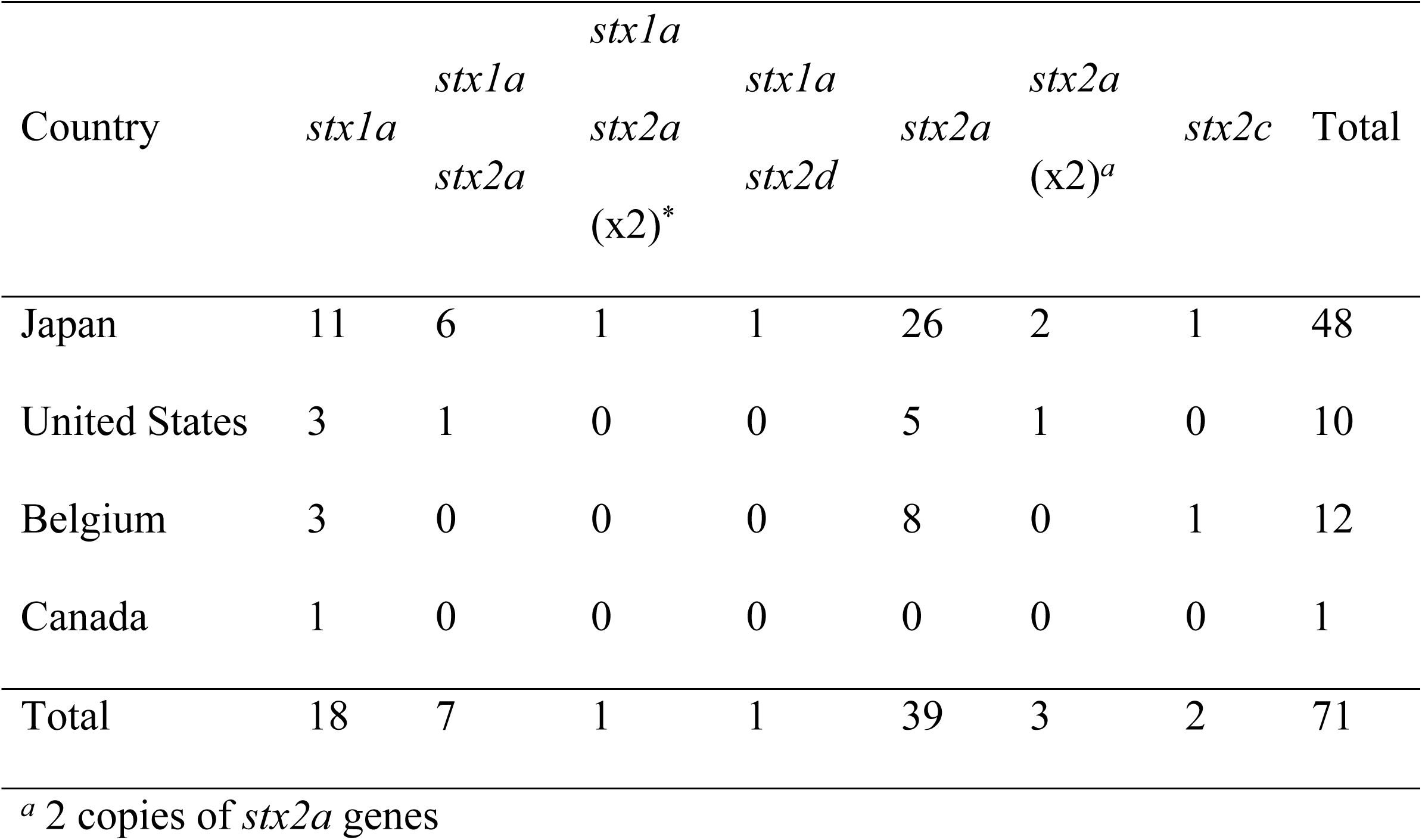
The *stx* genotypes of the O145:H28 strain analyzed in this study.

As the genome sequences of the Stx phages of 27 strains were already known (n=38), we determined those of the Stx phages in the remaining 43 strains (n=45). The Stx1a phage in an *stx1a*/*stx2d*-positive strain (strain 112808, whose Stx2d phage was previously sequenced but Stx1a phage was not) was also sequenced to obtain the full set of genome sequences of Stx phages (n=84) and determine their integration sites (Fig. 1, Table S1).

Of the 84 phages, 33 were S-Stx2a phages, and 21 were L-Stx2a phages. The remaining 30 phages encoded other types of Stxs, and all were L-phages (27 Stx1a, two Stx2c, and one Stx2d). For the integration sites, five loci (*argW*, *wrbA*, *attB* in the *ompW* prophage, *yecE*, and *sbcB*) were identified. The L-Stx2a phage was duplicated in strain 112648 and integrated into *attB* in the *ompW* prophage (referred to as *attB*_in_PP*ompW*) and the *yecE* loci [36]; thus, these phages were considered one L-Stx2a phage. Although three strains (12E129, RM12367-C1, and H27V05) contained two Stx2a phages, they were included as different phages because they showed considerable sequence variation (Fig. S1); thus, a total of 83 Stx phages were analyzed in subsequent analyses.

### Dynamics of Stx phages in O145:H28 isolates

To analyze the sequence similarities of the 83 Stx phages, we performed all-to-all sequence comparisons using the Mash program [37] and constructed a dendrogram using the complete linkage method based on pairwise Mash distances. The 83 phages were clearly divided into L-phages and S-phages (Fig. 2), but multiple phage clusters were detected with a threshold of 0.05 in both types of phages (PC1-PC5 in L-phages and PC6 and PC7 in S-phages).

**Fig. 2.**
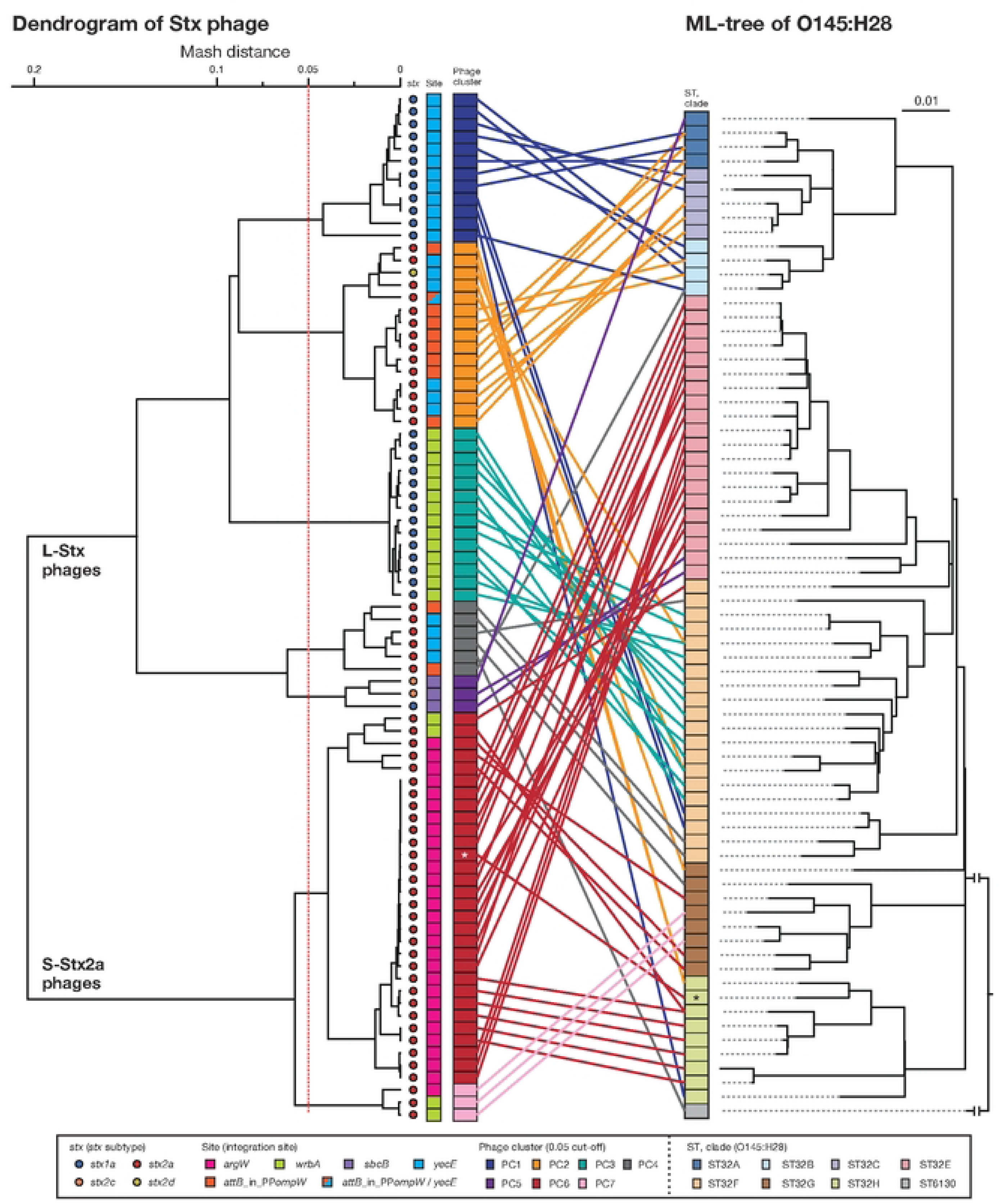
Sequence similarities among Stx phages found in the 71 O145:H28 strains. A dendrogram based on the Mash distance matrix of 83 Stx phage genomes is shown in the left panel, along with their *stx* genotypes, integration sites, and phage clusters, which were defined based on the pairwise Mash distance with a cutoff distance of 0.05. The phage indicated by *attB*_in_PP*ompW*/*yecE* was the duplicated L-Stx2a phages that were integrated into the *attB*_in_PP*ompW* and *yecE* loci in strain 112648. These duplicated phages were treated as one phage. The tree in the right panel is the same ML tree of O145:H28 strains shown in Fig. 1. Stx phages were connected to their host strains by lines colored according to the phage clusters. Strain Ech14022 and its S-Stx2a phage are indicated by asterisks.

In five phage clusters (PC1, 3, 4, 6, and 7), the encoded Stx was the same subtype. However, variations in Stx subtype were found in PC2 and PC5 (12 Stx2a and one Stx2d in PC2 and one Stx1a and two Stx2c in PC5), suggesting the replacement of *stx* in these two phage clusters. While the Stx1a phages at *yecE* and *wrbA*, and the Stx1a and Stx2c phages at *sbcB* formed distinct clusters (PC1, PC3, and PC5, respectively), the L-Stx2a and S-Stx2a phages were separated into two clusters (Fig. 2). Interestingly, although the L-Stx2a phages were separated into PC2 and PC4, both included phages at *attB*_in_PP*ompW* and *yecE*. Similarly, S-Stx2a phages were separated into PC6 and PC7, but both included phages at *argW* and *wrbA*. Variations in PC2 and PC4 can be easily generated because the *attB* sequences in *attB*_in_PP*ompW* and *yecE* are essentially the same [36]. In contrast, the variation in PC2 and PC4 was based on replacement of the integrase gene.

The within-cluster heterogeneity of Stx phages was more evident when the phylogeny of their host strains was considered (Fig. 2). While two clusters (PC3 and PC7) included phages found in the same host clade, the remaining five clusters included Stx phages found in multiple host clades. For example, PC1 phages were found in strains belonging to clades A, B, C, F, and H, and PC2 phages were found in clades A, B, C, F, G, and H. This finding indicates dynamic changes in Stx phages in each clade. The most striking case was the S-Stx2a phage of strain Ech14022 of clade H (indicated by an asterisk in Fig. 2), which had a nearly identical sequence to the S-Stx2a phages of clade E strains, suggesting recent interclade transfer of this phage.

### Stx2 production was greater in the strains carrying S-Stx2 phages than in the strains carrying L-Stx2 phages

To examine the variation in the level of mitomycin C (MMC)-induced Stx2 production across O145:H28 strains, we measured the Stx2 concentrations in the cell lysates of 45 *stx2*-positive strains available in our laboratory (29, 13, 2, and 1 strains carried S-Stx2a, L-Stx2a, L-Stx2c, and L-Stx2d phages, respectively), which covered seven of the eight clades in ST32. As shown in Fig. 1 and Table S1, the Stx2 production levels were highly variable between the strains (0.06-28.6 µg/ml). Moreover, the comparison of strains carrying S-Stx2a phages and those carrying L-Stx2 phages (including Stx2a, Stx2c, and Stx2d phages) revealed that the former strains produced significantly more Stx2 than the latter strains (19.8 vs. 4.1 µg/ml on average; *P* < 0.0001) (Fig. 3A). The Stx2 production level of strain H27V05, which carried two S-Stx2a phages, was average (19.5 µg/ml) among the S-Stx2a phage-carrying strains. However, the two strains (112648 and 12E129) that carried two L-Stx2a phages produced greater amounts of Stx2 (12.6 µg/ml and 15.0 µg/ml, respectively) than the other strains carrying L-Stx2 phages.

**Fig. 3.**
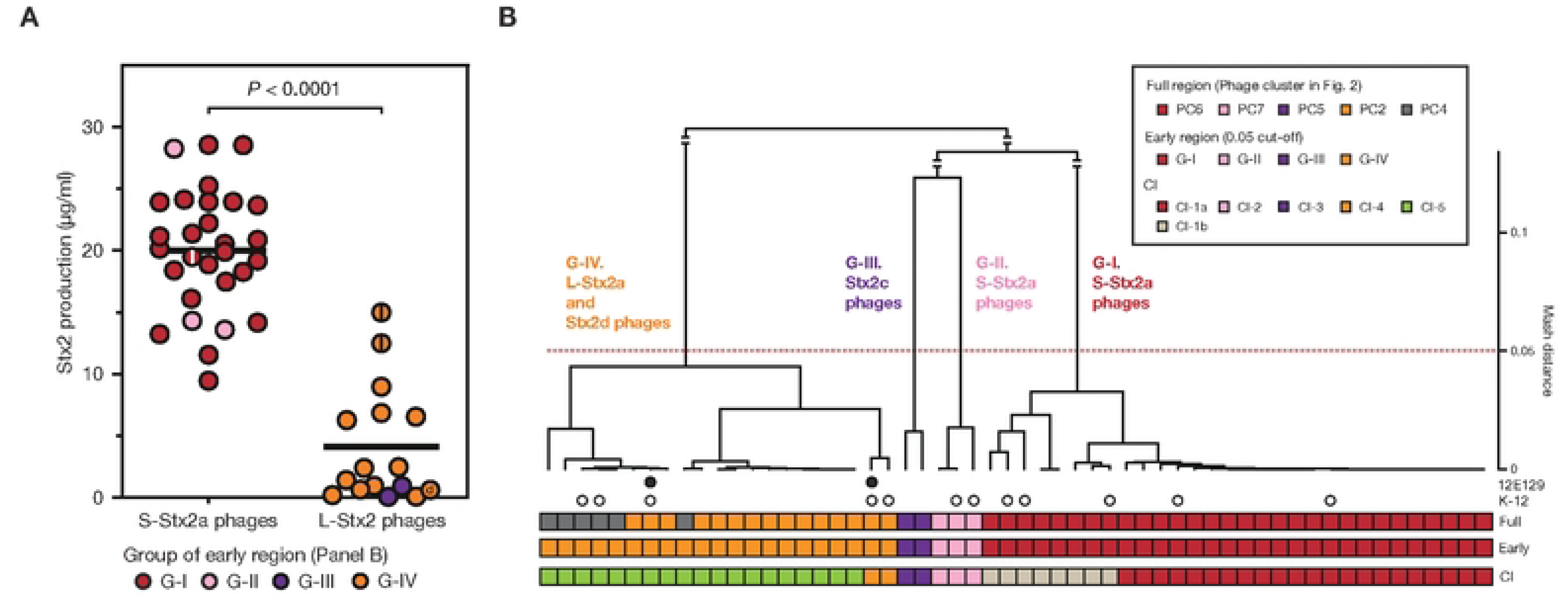
Stx2 production levels of O145:H28 strains harboring S-Stx2a phages and those harboring L-phages encoding Stx2a, Stx2c, or Stx2d and comparisons of early regions between Stx2 phage genomes. (A) Comparison of the Stx2 production levels between strains carrying S-Stx2a phages and those carrying L-Stx2 phages. Most L-phages encoded Stx2a, but two encoded Stx2c and one encoded Stx2d. The Stx2 production level of each strain is presented as the mean value of biological triplicates. Each strain is colored according to the groups defined based on the sequence similarity of the early region of their Stx2 phages in Fig. 3B. Strains carrying two Stx2a phages are indicated by the center lines in circles. The Stx2d phage-carrying strain is indicated by “d”. (B) Sequence similarities of the early regions of Stx2 phages. A dendrogram was constructed based on pairwise Mash distances. Stx2 phages were divided into four groups (threshold: 0.05). Phage clusters, which were defined based on the sequence similarity of full-length genomes (shown in Fig. 2), and the types of CI repressors are also indicated. Two L-Stx2a phages in strain 12E129 are indicated by filled circles. Stx2a phages lysogenized into K-12 (see Fig. 4) are indicated by open circles.

### Variations in the early genes of S– and L-Stx2 phages

Stx2 production levels were clearly different between the strains harboring S– and L-phages. However, as mentioned before, Stx2 production is tightly coupled with phage induction, which is achieved by the expression of early genes and the activation of late gene promoters [26,27]. Therefore, to examine the relationship between the variation in Stx2 phages and the Stx2 production levels in host strains in more detail, we performed an additional clustering analysis of the Stx2 phages (n=56; the Stx2 production levels of their host strains were determined) based on the pairwise Mash distances of their early regions (Fig. 3B). These phages were classified into four groups (referred to as G-I, G-II, G-III, and G-IV) with a threshold of 0.05, the same threshold that is used for the analysis of full-length phage genomes. This grouping correlated well with that based on full-length phage genomes; G-I contained all S-Stx2a phages in PC6, G-II contained all S-Stx2a phages in PC7, and G-III contained both L-Stx2c phages in PC5. However, all L-Stx2a phages and one L-Stx2d phage in PC2 and PC4 were grouped together into G-IV, indicating that they contained similar early regions, which were distinct from those in the S-Stx2a and L-Stx2c phages.

We further analyzed the sequence variation in CI repressors and Q anti-terminators, which are key regulators of phage induction [38], among the 56 Stx2 phages. In this analysis, we included the CI and Q proteins of the lambda, Sp5, and 933W phages. Sp5 and 933W, which were found in the O157:H7 strains Sakai and 933W, respectively [13,27], were representative S-Stx2a phages. Note that while the Q proteins of Sp5 and 933W were identical, their CI proteins were very different (19.1% identity).

The Q proteins of the S-Stx2a, L-Stx2c, and L-Stx2a phages were conserved (identical in sequence) in each type of Stx phage, except for a unique L-Stx2a phage of strain EH1910, but the Q proteins differed from each other type of Stx phage (Fig. S2). The Q protein of the L-Stx2d phage was also identical to the Q proteins of L-Stx2a phages. Among the four types of Q proteins, those of the S-Stx2a phages were most similar to those of Sp5 and 933W (91% identity).

The CI proteins of the Stx2 phages of O145:H28 strains were divided into several types with distinct sequences (referred to as CI-1, CI-2, CI-3, CI-4, and CI-5) (Figs. 3B and S3). The CI-1 type was further divided into two subtypes (CI-1a and CI-1b; 68.6% identity between the two subtypes). Among the six types of CI proteins, CI-2 was identical to the CI of 933W. Although the CI types correlated well with the phage groups defined by the sequence similarities of the early regions (Fig. 3B), S-Stx2a phages belonging to G-I were divided into to the CI-1a and CI-1b subtypes. G-IV phages (including all L-Stx2a phages and one L-Stx2d phage) were also divided into two types: CI-4 (comprising two L-Stx2a phages) and CI-5 (comprising the other G-IV phage). Notably, the CI proteins of the two L-Stx2a phages in strain 12E129 (both in G-IV) were CI-4 and CI-5.

These results indicated that although the Stx2 phages of O145:H28 strains were classified into S– and L-phages, both were further divided into several groups/types based on the sequences of the early genes and the Q and CI proteins, which may have some impact on phage induction. Notably, all S-Stx2a phages belonging to group G-II shared identical CI-2 and Q proteins with 933W, suggesting that the induction process of this group of S-Stx2a phages may be similar to that of 933W.

### Variation in phage induction patterns between different types of Stx2a phages under the K-12 background

To understand how the variation in the early regions of Stx2a phages affects their phage induction and associated Stx2 production, we generated K-12 lysogens of Stx2a phages belonging to the G-I, G-II or G-IV groups with various CI types (Table S2) and obtained lysogens from 12 phages (Fig. 4A and Table S2). Focusing on the Stx2a phage, we did not generate lysogens of the Stx2c and Stx2d phages. Analyses of these lysogens revealed notable variations in MMC-induced phage induction and Stx2 production, which were associated with phage type. Hereafter, Stx2a phages are referred to as Stx2a_xxx (where xxx denotes the O145:H28 host strain; i.e., Stx2a_112648).

**Fig. 4.**
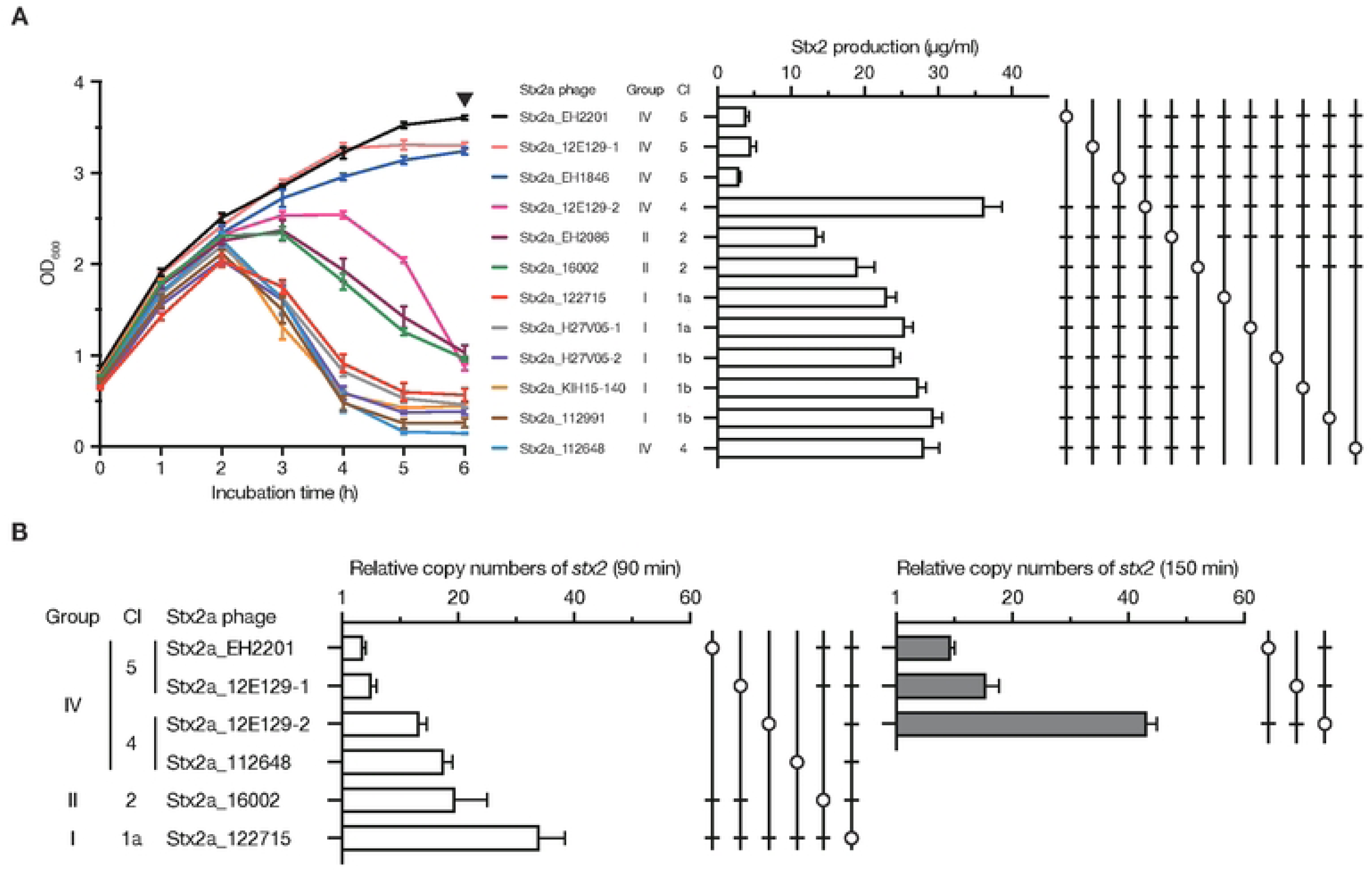
Analysis of 12 *E. coli* K-12 lysogens carrying different types of Stx2a phages. (A) Lysis curves (left panel) and Stx2 production levels (right panel) of MMC-treated lysogens. Stx2 production levels are presented as the Stx2 concentrations in the cell lysates obtained after 6 h of MMC treatment (mean values with standard errors of biological triplicates are shown). The samples indicated by open circles were significantly different (*P <* 0.05) from the samples marked by bars. (B) Relative copy numbers of *stx2a* in the cellular DNA samples of each lysogen at 90 min or 150 min after the start of MMC treatment. Relative copy numbers were determined by calculating the ratio of the copy number of *stx2a* relative to that of the *rluF* gene. The mean values with standard errors of biological triplicates are shown. Statistically significant differences between samples are also shown in the same way as those in panel A.

#### Lysis pattern

We observed rapid and complete lysis in all lysogens of G-I S-Stx2a phages, independent of their CI-1 subtypes (1a or 1b), although those of G-II S-Stx2a phages (CI-2) showed delayed lysis (Fig. 4A). The lysogens of G-IV L-Stx2a phages (CI-4 or CI-5) showed variable lysis patterns. While the Stx2a_112648 (CI-4) lysogen showed rapid and complete lysis, the Stx2a_12E129-2 (CI-4, the second L-Stx2a phage of strain 12E129) lysogen showed delayed lysis similar to but more prominent than G-II phage lysogens. Moreover, no detectable lysis was detected for the lysogens of CI-5-type phages, including Stx2a_12E129-1 (one of the two L-Stx2a phages of this strain).

#### Stx2 production level

Stx2 concentrations in the cell lysates of all G-I S-Stx2a phage lysogens were greater than 20 µg/ml (Fig. 4A). Compared with these lysogens, the G-II S-Stx2a phage lysogens produced lower amounts of Stx2, particularly the Stx2a_EH2086 lysogen. Among the lysogens of the five G-IV L-Stx2a phages, while the lysogens of CI-5 phages exhibited remarkably low Stx2 production, those of CI-4 phages produced large amounts of Stx2 comparable to or even greater than those of G-I S-Stx2a phages.

#### Phage induction as measured by the *stx2a* gene copy number

To examine the induction efficiency of the G-IV L-Stx2a phages that showed variable lysis patterns and Stx2 production, we determined the copy numbers of *stx2* in the cellular DNA of four G-IV L-Stx2a lysogens (two CI-4 and two CI-5 phages). Lysogens of two S-Stx2a phages (G-I and G-II) were also analyzed for comparison (Fig. 4B). At 90 min after the start of MMC treatment, the copy number of *stx2* relative to that of a housekeeping gene (*rluF)* increased in all phages (see Table S2 for more details), but there were notable variations; the copy numbers of the two G-IV/CI-5 phages were lower than those of the other phages. At 150 min, the copy numbers increased in all three G-IV phages examined (the other three were not examined because lysis started at this time point), but the copy numbers of both CI-5 phages were significantly lower than that of the CI-4 phage. These results indicated that G-IV/CI-5 phages had lower phage induction efficiency, which explains the lower Stx2 production by their K-12 lysogens.

Taken together, these analyses revealed that, of the two types of G-IV L-Stx2a phages, lysogens of the CI-5 type, which represent the major CI type in the L-Stx2a phages of O145:H28, produced lower amounts of Stx2 due to their lower phage induction efficiency. In contrast, the CI-4 type is efficiently induced and confers much greater Stx2 production by host strains, which is comparable to that of G-I S-Stx2a phages.

### Comparison of the early regions between different types of Stx2a phages

To examine the relationship between variations in the Stx2a phage genomes and the observed phenotypes of their lysogens, we compared the genetic structures of the early regions between the 12 Stx2a phages whose K-12 lysogens were analyzed (Fig. 5). Consistent with the data shown in Fig. 3B, the early regions exhibited overall sequence similarities within each group/CI type. However, in addition to the *cI* genes in G-I (CI-1a or CI-1b) and G-IV (CI-4 or CI-5), several genes exhibited notable within-group variations, which may also be linked to the observed within-group/CI-type differences in phage induction and Stx2 production by lysogens (Fig. 4). For example, replacement of a 2.9-kb segment encoding *cI* and three additional genes occurred between the CI-4 and CI-5 types of G-IV/L-Stx2a phages, which showed different phage induction patterns.

**Fig. 5.**
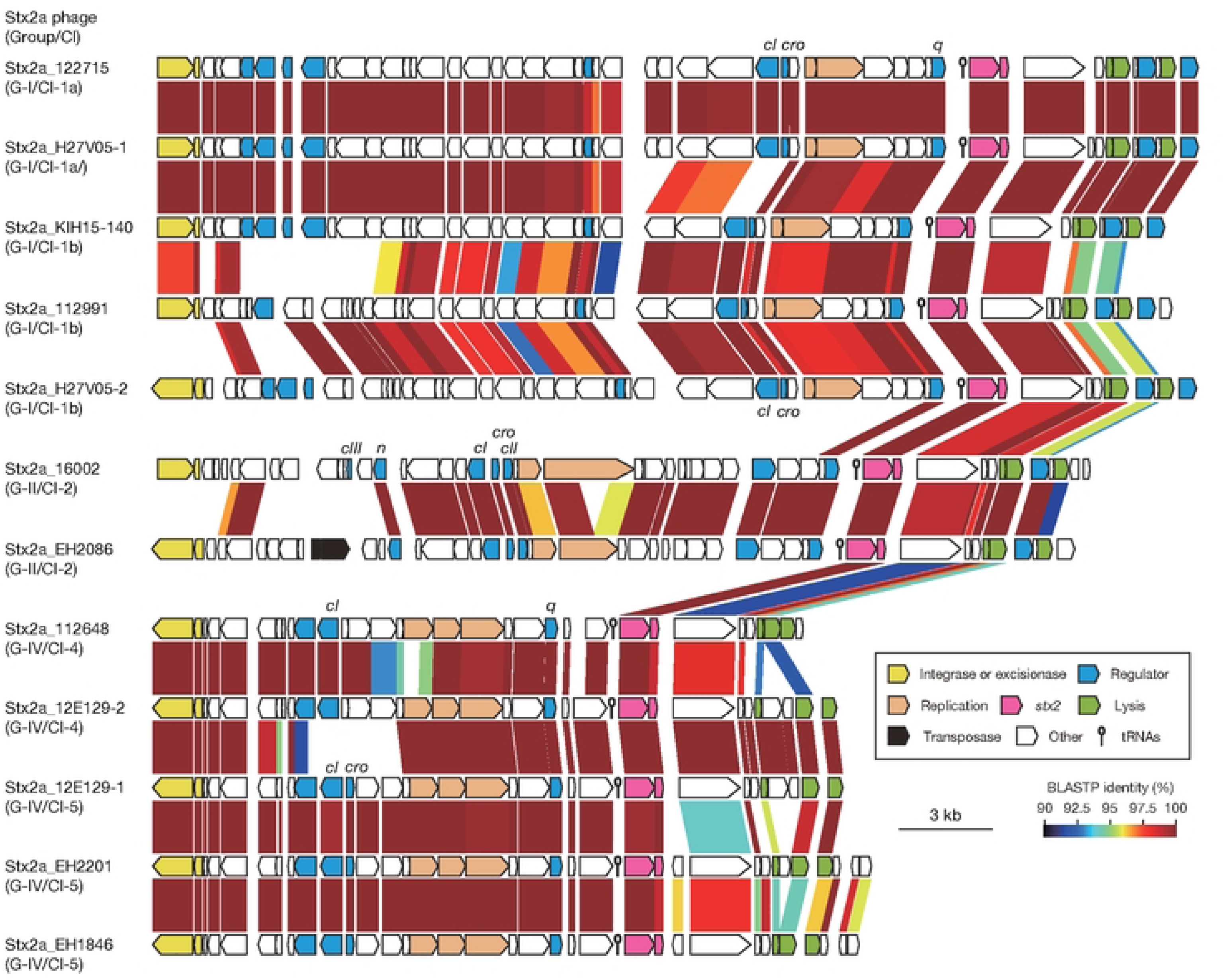
Comparison of the early regions of 12 Stx2a phages lysogenized into K-12. The genetic structures of the early regions of seven S-Stx2a phages (two G-I/CI-1a, three G-I/CI-1b, and two G-II/CI-2 phages) and five L-Stx2a phages (two G-IV/CI-4 and three G-IV/CI-5 phages) are drawn to scale. Amino acid sequence homologies are shown by shading with a heatmap.

### Effects of lysogenization by two L-Stx2a phages on the Stx2 production level

From the three strains that carried two Stx2a phages, we selected strains 112648 and 12E129 and generated mutants lacking each of the two L-Stx2a phages (Fig. 6A). Using these mutants and wild-type (WT) strains, we analyzed the effects of lysogenization of the two L-Stx2a phages on Stx2 production upon MMC-induced phage induction (Fig. 6B).

**Fig. 6.**
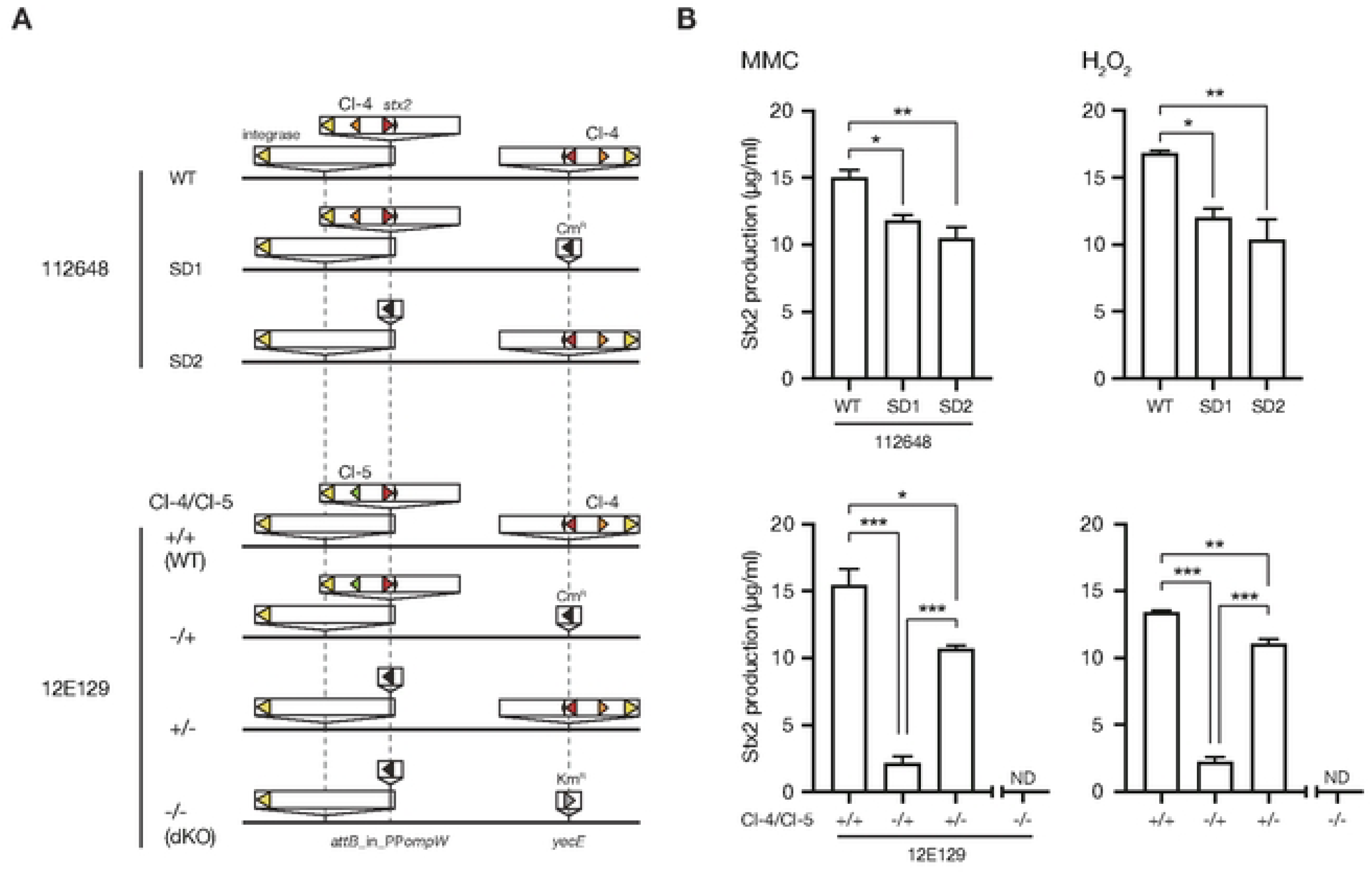
Stx2 production by O145:H28 strains harboring two L-Stx2a phages and their L-Stx2a phage-deletion mutants. (A) Schematic representation of the genomic locations of L-Stx2a phages in the wild-type (WT) strains and their Stx2a phage deletion mutants. Strain 112648 carried two identical CI-type L-Stx2a phages at the *attB*_in_PP*ompW* and *yecE* loci, and strain 12E129 carried a CI-type phage and a CI-4-type phage at these loci. Stx2a phage-deletion mutants were generated by replacing either one or both Stx2a phages with antimicrobial resistance gene cassettes as indicated. (B) Stx2 production by the WT strains and their Stx2a phage deletion mutants upon MMC or H_2_O_2_ treatment is shown. The Stx2 production levels are presented as the Stx2 concentrations in the cell lysates obtained after 6 h of treatment with MMC or H_2_O_2_ (the mean values with standard errors of biological triplicates are shown). Statistically significant differences are marked by asterisks (*, *P* < 0.05; **, *P* < 0.01; ***, *P* < 0.001). ND, not detected.

#### Lysogenization of two duplicated copies of the L-Stx2a phage

Strain 112648 carried two duplicated copies of G-IV/CI-4 phages (Fig. 6A). Mutants lacking either copy (SD1 or SD2) produced similar amounts of Stx2, which was significantly lower than that of the WT. This result suggested that the effect of duplication of this L-Stx2a phage on Stx2 production was additive.

#### Lysogenization of two L-Stx2a phages of different CI types

Strain 12E129 carried two types of G-IV L-Stx2a phages (CI-5 and CI-4 types named Stx2a_12E129-1 and Stx2a_12E129-2, respectively). Mutants carrying only one of the two phages (CI-4(-)/CI-5(+) and CI-4(+)/CI-5(-) in Fig. 6) produced significantly less Stx2 than the WT. In particular, deletion of the CI-4 phage drastically reduced Stx2 production. Thus, the impact of CI-4 phage lysogenization on Stx2 production was much greater than that of CI-5 phage lysogenization. This result is consistent with the abovementioned difference between the two types of L-Stx2a phages (Fig. 4).

### Differences in virulence of the lysogens of the two types of L-Stx2a phages

Generation of the mutants of strain 12E129, which carried either the CI-4 or CI-5 L-Stx2a phage, allowed us to evaluate the contribution of the two types of L-Stx2a phages to the virulence of this strain. Before performing *in vivo* experiments, we confirmed that the difference in Stx2 production between the WT strain and mutants observed upon MMC treatment was reproducible upon treatment with hydrogen peroxide (H_2_O_2_), a molecule that can be produced by neutrophils in the intestine during STEC infection [39] (Fig. 6B; note that this was also the case for the 112648 strain and its mutants). Then, using germ-free mice treated with dextran sulfate sodium (DSS), which induces colitis [40], we evaluated the virulence of the WT 12E129 strain and its mutants carrying either CI-4 or CI-5 L-Stx2a phage. A 12E129 mutant lacking both phages was also generated and used as a negative control after confirming that it produced no Stx2 (dKO in Figs. 6 and 7). Inoculated bacteria were stably colonized on day 5, as was determined by counting the WT and mutant cells in mouse feces (over 10^9^ CFUs/g feces; Fig. S4). Although some differences in CFU count were observed among the strains/mutants on day 5, no significant differences were detected on day 8.

**Fig. 7.**
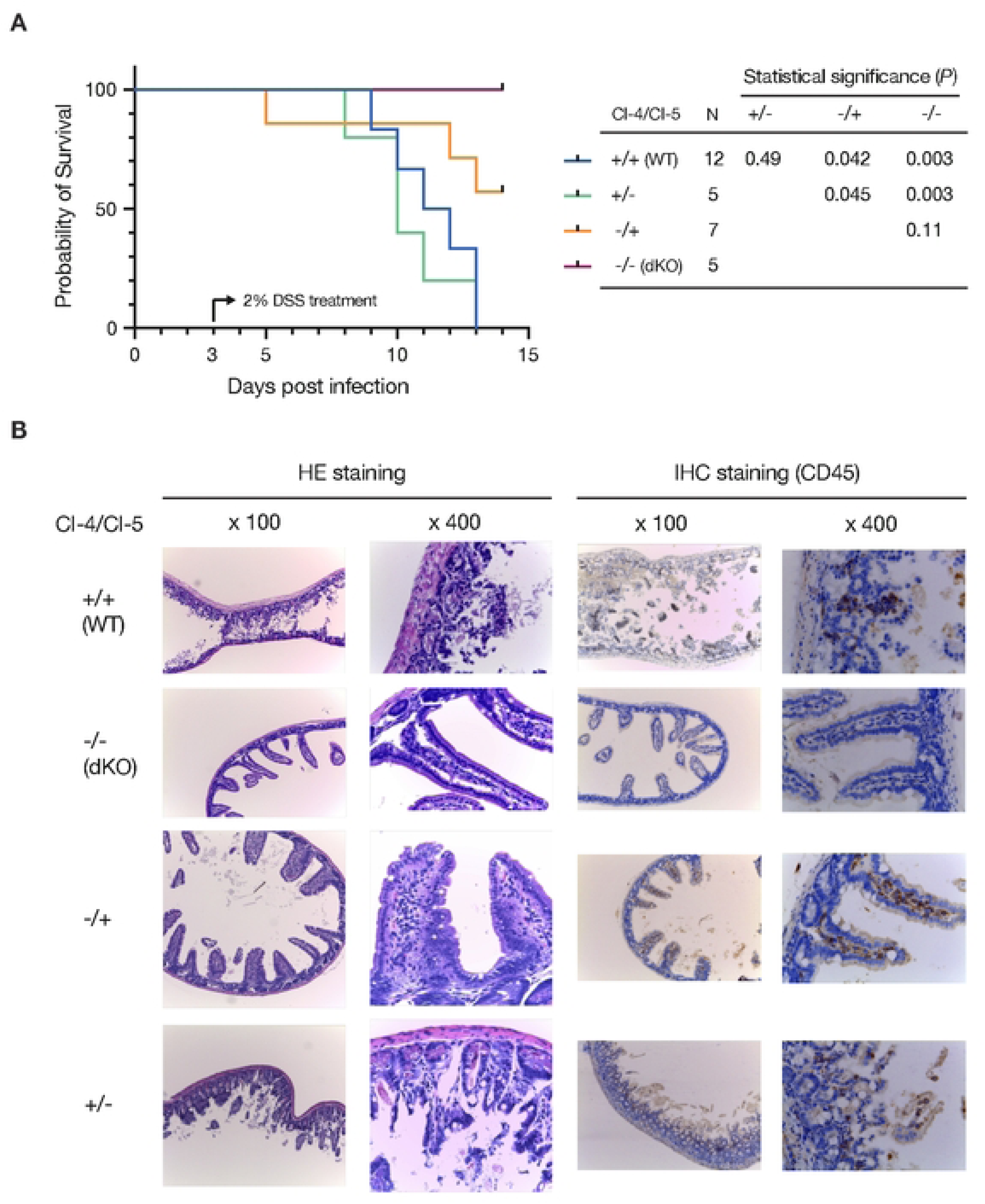
Evaluation of the virulence of strain 12E129 with two different types of L-Stx2a phages and Stx2a phage deletion mutants. The virulence of strain 12E129 and its Stx2a phage-deletion mutants in germ-free mice was evaluated using a DSS-induced colitis model. Strain 12E129 carried one CI-4 type phage and one CI-5 type phage. Each mutant carried either one or neither of the two L-Stx2a phages, as shown in Fig. 6. (A) Kaplan-Meier survival curves of mice inoculated with strain 12E129 and its Stx2a phage deletion mutants are shown in the left panel. Beginning on day 3, 2% DSS-containing water was given to the mice until the end of the experimental period. In the right panel, the significance of the differences between mouse groups calculated by the log-rank test and the number of mice tested are shown. (B) Ileal tissue sections of mice inoculated with strain 12E129 and its Stx2a phage-deletion mutants are shown. The sections were analyzed by hematoxylin-eosin (HE) staining and immunohistochemistry (IHC) staining of CD45.

Mice inoculated with the WT strain began to die on day 9, and all the mice died on day 13; however, no mice inoculated with the dKO mutants died within 14 days (Fig. 7A). Importantly, mice inoculated with the mutant carrying the CI-4 phage alone showed a survival pattern similar to that of WT-inoculated mice, and all the mice died by day 13. In contrast, more than half of the mice inoculated with the mutant carrying the CI-5 phage alone survived at the end of the experiment, and this survival rate significantly differed from that of WT-inoculated mice (and differed from that of mice inoculated with the mutant carrying the CI-4 phage). Consistent with these results, marked villus destruction and inflammatory cell infiltration were observed in the ileum of mice inoculated with the WT strain and the mutant strain harboring the CI-4 phage (Fig. 7B). In contrast, in the ileum of mice inoculated with the mutant strain carrying the CI-5 phage, the morphology of the villi was maintained, although mild inflammatory responses were observed. These results suggested that the CI-4 L-Stx2a phage confers increased virulence to host strains when compared to the CI-5 L-Stx2a phage.

### Distribution of L-Stx2 phages encoding CI-4 and CI-5 types of CI repressors in *E. coli*

A search in the NCBI database for Stx phages encoding CI repressors highly similar to the CI-4 or CI-5 type (>98% sequence identity) was used to identify four Stx phages (one L-Stx2a and three L-Stx1a phages) encoding a CI-4 type repressor in *E. coli* strains outside of the O145:H28 lineage (Table S3). Among these phages, the L-Stx2a phage of the O121:H19 strain exhibited high sequence similarity to the CI-4 L-Stx2a phage of O145:H28, but the three L-Stx1a phages shared only limited genes with the O145:H28 phage (Fig. S5). In contrast, Stx2 phages encoding CI-5-type CI repressors were identified in 30 strains belonging to 15 serotypes, including major STEC serotypes such as O157:H7, O26:H11, and O111:H8, and all were L-phages (25 Stx2a, one Stx2c, and seven Stx2d phages). These results suggest the rare distribution of CI-4-type L-Stx phages and the wide distribution of CI-5-type L-Stx phages in STEC strains.

## Discussion

Our systematic analysis of the full set of Stx phages in STEC strains covering the entire O145:H28 lineage revealed marked genomic diversity of Stx phages. Although the Stx phages were divided into long– and short-tailed phages (referred to as S– and L-phages, respectively, in this article) based on the types of late genes, their sequence similarity, as defined by pairwise Mash distance analysis, was used to classify them into seven phage clusters (PC1∼PC7; Fig. 2). While all Stx1 phages were L-Stx1a phages and belonged to PC1 or PC3, Stx2 phages included both L– and S-phages and were further classified into four groups (G-I∼G-IV; Fig. 3) based on the sequence similarity of their early regions and six CI types based on the sequences of CI repressors (Fig. 3).

An important finding obtained by analysis of the intra-serotype diversity of Stx phages was that Stx phages belonging to the same cluster were distributed in multiple clades of O145:H28 (Fig. 2). Although a possible case of direct interclade transmission of Stx2a phage was found in PC6, the variations observed for the entire lineage of O145:H28 strains suggest that Stx phages belonging to each cluster are widely circulating with continuous diversification in sequence. Sequence diversification within each cluster included the replacement of genomic segments, leading to within-cluster variations in integration sites (*argW* or *wrbA* for the S-Stx2 phages in PC6 and PC7), Stx types (the emergence of an Stx2d phage from the L-Stx2a phages in PC2 and the presence of Stx2c and Stx1a phages in PC5), and CI types (CI-4 or CI-5 type of L-Stx2a phages in PC2). Wide circulation of each phage cluster likely allowed for the acquisition of similar phages by host strains with various phylogenetic backgrounds. Repeated acquisition of Stx phages may induce the loss of resident Stx phages, leading to within-host clade variation in Stx phages. Similar changes/variations were observed for Stx1a phages in the ST21 lineage of STEC O26:H11 [25] and Stx2a phages in clade 8 of STEC O157:H7 [30]. However, notably, different distribution patterns of Stx phages were also observed in STEC O121:H19 and the STEC belonging to clonal complex 119 (CC119). In these STECs, systematic analyses of Stx phages have been conducted and revealed the stable maintenance of an S-Stx2a phage and an L-Stx2a phage in the major lineage of each STEC, respectively [29,31].

From a medical point of view, the more important findings of this study were the variations in Stx2 production levels by host strains, which were linked to the variations in Stx2 phages, because Stx2 is associated with the severity of STEC infection [6,41,42]. We first observed a striking difference in Stx2 production between the strains carrying S-Stx2a phages (G-I and G-II) and those carrying L-Stx2 phages (L-Stx2c phages in G-III and L-Stx2a phages in G-IV): the former strains produced more Stx2 than the latter strains, with only a few exceptions (Fig. 3A). Analysis of K-12 lysogens of 12 Stx2 phages belonging to different groups/CI types (Fig. 4A) confirmed that S-Stx2 phages confer greater Stx2 production by host strains. This analysis further revealed that L-Stx2a phages encoding the CI-4 type CI repressor confer increased Stx2 production, comparable to that of S-Stx2a phages, to host strains (Figs. 4 and 5). Moreover, we confirmed that the CI-4 type L-Stx2a phage is responsible for the increased virulence of the host strain in the O145:H28 background (Fig. 6). Similar findings related to the association of increased Stx2 production in host strains with Stx2a phages with specific types of early regions have been reported for the S-Stx2 phages of O157:H7 STEC [28,30]. Thus, the present study further highlighted the importance of analyzing the variation in Stx2 phage genomes in the surveillance of STEC. In this regard, although distinguishing S-Stx2a and L-Stx2a phages is important, we should also consider the presence of CI-4-type L-Stx2a phages, which are associated with increased Stx2 production and increased virulence, although this type of Stx2 phage appears to be rarely distributed in STEC.

Regarding the mechanism underlying the variation in the Stx2 production by STEC strains conferred by different types of Stx2 phages, our analysis of K-12 lysogens revealed that the increased induction efficiency of S-Stx2a phages and CI-4-type L-Stx2a phages contributed to increased Stx2 production by the host strains (Fig. 4). The observed differences in the induction efficiency of Stx2 phages are linked to differences in the early region of the phages, including differences in CI repressors (Fig. 5). However, further analyses are required to understand at the molecular level how differences in the early region are involved in determining the efficiency of phage induction and the levels of Stx2 production by host strains.

## Materials and Methods

### Bacterial strains

The initial O145:H28 strain set included 59 strains that were available in our laboratory and sequenced in our previous study [33] and 18 strains with complete genome sequences (the plasmid genome was not available for strain 2015C-3125). The genome sequences of the 18 previously reported strains [34,35,43,44] were downloaded from NCBI. Six of the 18 strains were excluded from the strain set because their recombination-free core sequences were identical to those of other strains (see Table S1 for the strains included in the final set). Therefore, in this study, 71 strains were analyzed.

### Determination of complete genome sequences

The genomic DNA of strains 16003 and 12E115 was purified using Genomic-tip 100/G (Qiagen). Libraries for Illumina sequencing (average insert size: 700 bp) were prepared using the NEBNext Ultra II FS DNA Library Prep Kit (New England Biolabs) and sequenced using Illumina MiSeq to generate 300 bp paired-end reads. These genomes were additionally sequenced using MinION with R9.4.1 flow cells (Nanopore) for 68 (16003) or 96 h (12E115). Nanopore reads were trimmed and filtered as described previously [29] and assembled along with the Illumina reads of each strain using Unicycler v0.4.8 [45] to obtain the finished genome sequences. The sequences of the Illumina and Nanopore reads and the complete genome sequences of these strains were deposited in DDBJ/EMBL/GenBank under the BioProject accession numbers starting from PRJDB8147 (see Table S1 for each accession number).

### Phylogenetic analysis

Phylogenetic analysis of the initial strain set (n=77) was performed based on the SNPs identified on the prophage/integrative element/IS-free and recombination-free chromosome backbone that was conserved across all genomes by Gubbins [46] and MUMmer [47] using the genome of strain 10942 as a reference. A maximum likelihood (ML) tree was constructed with RAxML [48] as previously described [36] and displayed using FigTree v1.4.4 (http://tree.bio.ed.ac.uk/software/figtree/).

### Analyses of integration sites and the sequencing of Stx phages

Stx phages integrated into the *attB*_in_PP*ompW* and *yecE* loci in 54 O145:H28 draft genomes were previously described [36]. Stx phages integrated into *argW*, *wrbA*, and *sbcB* in these 54 genomes were identified by the same strategy as that we previously employed (schematically shown in Fig. S6; see Table S4 for the primers used). The entire genome sequence of each phage was determined by sequencing long PCR products as previously described [36]. DFAST [49] and GenomeMatcher (v3.0.2) [50] were used for annotating phage genomes and comparing Stx phage genomes, respectively.

### Clustering analyses of Stx phage genomes and typing of CI and Q proteins

All-to-all phage genome comparison was performed for the three sets of genomes (the entire genomes of 83 Stx phages, the entire genomes of 54 Stx2 phages, and the early regions of 54 Stx2 phages) using Mash v2.0 [37] with default parameters to generate pairwise Mash distance matrices. Based on each matrix, Stx phages and the early regions of Stx phages were clustered with a cutoff of 0.05 as previously described [51]. The amino acid sequences of CI and Q encoded by each Stx2 phage were aligned with the proteins from phages lambda, Sp5, and 933W by MUSCLE in MEGA v10.1.8 [52], and dendrograms were generated based on these alignments using the UPGMA algorithm in MEGA v10.1.8.

### Search for Stx phages encoding CI-4– or CI-5-type CI repressors in a public database

CI-4– or CI-5-type CI repressors were identified in the nucleotide collection (nr/nt) in the NCBI database (last access: January 29, 2024) by TBLASTN using the amino acid sequences of the CI-4-type repressor in phage Stx2a_12E129-2 and the CI-5-type repressor in phage Stx2a_12E129-1 as queries with a threshold of >98% identity and 100% coverage. After excluding the sequences in *E. coli* O145:H28 strains, those on apparently full-length prophage genomes were selected. The *stx* subtypes of these prophages were determined by BLASTN as previously described [33].

### Lysogenization of Stx2a phages into *E. coli* K-12

Prophages were induced with MMC as described previously [36]. At 6 h after the start of MMC treatment, each culture was treated with 1/500 volume of chloroform and centrifuged at 12,000 ×g for 20 min at 4 °C. The supernatant was filtered through 0.22-µm pore size filters (Millipore). Phage particles in the supernatant were collected by polyethylene glycol/NaCl precipitation as described previously [53] when necessary. Serially diluted phage lysates (100 µl) were prepared with SM buffer [53], mixed with K-12 MG1655 cells suspended in 1 ml of lysogeny broth (LB) containing 10 mM CaCl_2_ (2.5-4.0 OD_600_), and incubated for 20 min at 37 °C. Then, the phage/bacterial cell mixture was mixed with 5 ml of top agar (0.75% w/v LB agar) containing 10 mM CaCl_2_ and 1.0 µg/ml MMC and spread onto bottom LB agar (1.5% w/v). After overnight incubation at 37 °C, the plaques were picked, cultured in LB overnight at 37 °C, and spread onto LB agar. Several colonies were randomly selected and analyzed by colony PCR using the *stx2*-specific primers stx2-F and stx2-R [54] to confirm Stx2a phage lysogeny. We tried to generate lysogens for the 17 Stx2a phages listed in Table S2 and succeeded in obtaining the lysogens of 12 phages. We purified genomic DNA from the 12 lysogens using the DNeasy Blood and Tissue Kit (Qiagen) and determined the genome sequences of Stx2a phages lysogenized in each lysogen by the long PCR-based strategy, as shown in Fig. S6.

### Deletion of Stx2 phage genomes

To delete the Stx2 phage genomes integrated into *attB*_in_PP*ompW* or *yecE* in strains 112648 and 12E129, we introduced the Red recombinase encoding plasmid, pKD46 [55], into these strains. A DNA fragment containing the chloramphenicol resistance (Cm^R^) cassette and terminal 55-nt extensions homologous to the *attL* and *attR* flanking regions of each locus was generated by the 2-step tailed PCR method using two sets of primers and pKD3 as a template. The 1.1-kbp PCR product was purified, treated with DpnI, and introduced into 112648 and 12E129 cells harboring pKD46, in which arabinose-inducible Red recombinase was expressed. The deletion of each Stx2a phage in the Cm^R^ transformants was confirmed by colony PCR using EmeraldAmp MAX PCR Master Mix (TaKaRa) and specific primers. To generate a mutant lacking both Stx2a phages from strain 12E129, we first deleted the Stx2a phage at *attB*_in_PP*ompW* and then deleted it at *yecE* by repeating the same procedure, except the DNA fragment containing the kanamycin resistance (Km^R^) cassette was generated using pKD4 as a template. The sequences of primers used for these experiments are listed in Table S5. We attempted to delete the Stx2 phage genomes integrated into *argW* or *wrbA* in strain H27V05 by the same strategy but were unable to obtain these mutants.

### Determination of Stx2 production levels

Cell lysates of all tested strains, mutants, and K-12 lysogens were prepared as described previously [29], except for the final concentration of MMC (Wako; 1.0 µg/ml). The MMC concentration and sampling time were optimized based on the results of exploratory analyses using seven O145:H28 strains (see Fig. S7 for details). The Stx2 concentration in the lysate of each O145:H28 strain was determined by sandwich ELISA as previously described [29]. As the ELISA kit (RIDASCREEN Verotoxin; R-Biopharm AG) became unavailable in Japan during this study, the Stx2 concentrations of the lysates of strains 112648 and 12E129, their Stx2a phage-deletion mutants, and the K-12 lysogens were determined by the homogeneous time-resolved fluorescence energy transfer (HTRF) assay that was recently developed for Stx2 quantification [56]. H_2_O_2_-induced Stx2 production was quantified by determining the Stx2 concentration in the cell lysates, which were prepared as described above except for treatment with H_2_O_2_ (Wako; 3 mM at the final concentration), via the HTRF assay.

### Determination of the copy number of *stx2*

Overnight cultures of K-12 lysogens were inoculated in 3 mL of LB at 0.1 OD_600_ and grown to mid-log phase at 37 °C with shaking. Then, MMC was added to the cultures at a final concentration of 1.0 μg/ml. After 90 min or 150 min of incubation, the bacterial cells were collected, and the total cellular DNA was purified using a DNeasy Blood and Tissue Kit. The copy number of *stx2* in each cellular DNA sample was determined by droplet digital PCR using the EvaGreen assay (Bio-Rad) with the *stx2*-specific primers described above. The copy number relative to that of the *rluF* gene was determined by dividing the copy number of *stx2* by that of *rluF*. The *rluF* gene was amplified with the primers rluF-S (5’-GCACGCGCATCATGAACGTTAG-3’) and rluF-R (5’-CGTCGGTTAAATCGCGCCATTC-3’).

### Virulence assay

From the point of view of animal welfare, we selected a minimum strain set for a mouse virulence assay based on data obtained in *in vitro* experiments to reduce the number of mice used in the virulence assay. The O145:H28 strain 12E129 and its Stx2a phage-deletion mutants were cultured overnight in Tryptic Soybean Broth at 37 °C with shaking. The cultures were diluted with distilled water (DW) to 1.0 OD_600_. One milliliter of each OD-adjusted culture was added to 100 ml of DW to prepare contaminated drinking water containing STEC at approximately 10^7^ CFU/ml. Male germ-free C57BL/6N mice (5 weeks old; Clea Japan, Inc.) were divided into four groups and kept separately in sterilized vinyl isolators with an irradiated chow diet, autoclaved cages, bedding, and DW. The mice were given contaminated water *ad libitum* for one day (day 0). On day 3, the contaminated water was replaced with 2.0% dextran sulfate sodium (DSS) to induce gut inflammation. The conditions of the mice were checked twice a day until day 14, and the survival rate was compared between the groups. Stool culture was performed on days 1, 2, 5 and 8 to monitor the colonization of bacteria. For this monitoring, fresh feces were suspended in saline to 0.1 mg/ml, and a 10-fold serial dilution was made. Then, 0.1 ml of appropriately diluted sample was spread onto MacConkey agar plates. After incubating at 37 °C for 48 h, colonies were counted to determine the number of bacteria in the feces. Autopsy was performed on the mice that died during the observation period, and the mice survived until day 14. The terminal ileum was collected, fixed in 10% neutralized formalin, and embedded in paraffin, and 3-µm tissue sections were prepared for hematoxylin-eosin staining and immunohistochemistry targeting CD45 to evaluate destruction of the ileal mucosa and leukocyte infiltration. CD45 was detected by using an IHCeasy CD45 Ready-To-Use Kit according to the manufacturer’s instructions (Cosmo Bio Co., Ltd., Tokyo, Japan).

## Statistical analyses

An unpaired *t* test was performed to compare Stx2 production levels between the O145:H28 strains carrying S-Stx2a phages and those carrying L-Stx2 phages using Prism 9 software (GraphPad Software). One-way analysis of variance (ANOVA) followed by the Tukey–Kramer multiple comparison test was performed using R v4.1.0 [57] to compare Stx2 production levels between K-12 lysogens and between strain 12E129 and its Stx2a phage-deletion mutants and the *stx2* copy numbers between K-12 lysogens. For the comparison of the survival rates between mice given strain 12E129 and those given mutant strains, a log-rank test was performed using StatFlex v6.0 (Artec Co., Ltd., Osaka, Japan). *P* < 0.05 was considered to indicate statistical significance.

## Ethics statement

Animal experiments were carried out in accordance with Japanese legislation (Act on Welfare and Management of Animals, 1973, revised in 2012) and guidelines under the jurisdiction of the Ministry of Education, Culture, Sports, Science and Technology, Japan (Fundamental Guidelines for Proper Conduct of Animal Experiment and Related Activities in Academic Research Institutions, 2006). The protocols for the animal experiments were approved by the Animal Care and Use Committee of Kagawa University (Approval Number; 17627-3). Animal care, housing, feeding, sampling, observation, and environmental enrichment were performed in accordance with the guidelines.

## Acknowledgments

This research was supported by AMED under Grant Number 21fk0108611h0501, 22fk0108611h050, and 23fk0108611h0503 to TH and a KAKENHI from the Japan Society for the Promotion of Science (18K07116) to KN. We thank M. Horiguchi and K. Ozaki for providing technical assistance.

## Author Contributions

Conceptualization: Keiji Nakamura, Tetsuya Hayashi

Data Curation: Keiji Nakamura

Formal Analysis: Keiji Nakamura, Haruyuki Nakayama-Imaohji, Munyeshyaka Emmanuel, Itsuki Taniguchi

Funding Acquisition: Keiji Nakamura, Tetsuya Hayashi

Investigation: Keiji Nakamura, Haruyuki Nakayama-Imaohji, Munyeshyaka Emmanuel, Yasuhiro Gotoh

Methodology: Keiji Nakamura, Tomomi Kuwahara, Tetsuya Hayashi

Project Administration: Keiji Nakamura, Tetsuya Hayashi

Resource: Junko Isobe, Keiko Kimata, Yukiko Igawa, Tomoko Kitahashi, Yohei Takahashi, Ryohei Nomoto, Kaori Iwabuchi, Yo Morimoto, Sunao Iyoda

Software: Itsuki Taniguchi

Supervision: Keiji Nakamura, Tetsuya Hayashi

Validation: Keiji Nakamura, Haruyuki Nakayama-Imaohji, Munyeshyaka Emmanuel

Visualization: Keiji Nakamura

Writing – Original Draft Preparation: Keiji Nakamura, Tomomi Kuwahara, Tetsuya Hayashi

Writing – Review & Editing: Keiji Nakamura, Tetsuya Hayashi

## Figure legends

**Fig. S1.** Sequence similarities of the two Stx2a phage genomes found in each of the four O145:H28 strains. Dot plot presentation showing sequence similarity (window size of 2 kb; >99% sequence identity) between two Stx2a phages in strains 112648, 12E129, RM12367-C1, and H27V05. The integration site and genome size of each phage are indicated on the X– and Y-axes. Note that the two Stx2a phages in strain 112648 are completely identical.

**Fig. S2.** Variation in the sequences of Q antiterminator proteins encoded by the Stx2 phages of the O145:H28 strains. (A) The Q types of each Stx2 phage, defined based on the sequence similarity of Q proteins, are shown under the dendrogram constructed based on the pairwise Mash distances of early regions (the same dendrogram in Fig. 3). The Q protein of the L-Stx2a phage that contained a frameshift mutation is indicated by an asterisk. (B) Amino acid sequence comparison of four types of Q proteins found in the Stx2 phages of O145:H28 strains. A UPGMA tree was generated based on the alignment of Q sequences. The Q proteins of the lambda phage (No. NP_040642.1), Sp5 (No. BAA94139.1), and 933W (No. NP_049499.1) were included as references.

**Fig. S3.** A UPGMA tree generated based on the sequence alignment of CI repressor proteins. Although there were four variants in the CI-5 type of CI repressors (>97.8% identity), they are depicted together in this figure. The CI proteins of the lambda phage (No. NP_040628.1), Sp5 (No. BAA94122.1), and 933W (No. NP_049485.1) were included as references.

**Fig. S4.** Colonization of strain 12E129 and its Stx2a phage-deletion mutants in the intestines of germ-free mice. The mean CFUs per gram of feces are shown with standard errors. One mouse inoculated with the mutant carrying the CI-5 L-Stx2a phage alone (indicated by “-/+”) accidentally died on day 5, and this mouse was excluded from the analysis. Two fecal samples could not be obtained (the day 5 sample of a mouse infected with the wild-type (WT) 12E129 strain and the day 8 sample of a mouse infected with the WT strain); these samples were also not included in this analysis. The levels of significance are indicated by asterisks (*, *P* < 0.05; **, *P* < 0.01).

**Fig. S5.** Comparison of the Stx genomes encoding CI-4-type CI repressors. The genomic organization of three L-Stx2a and three L-Stx1a phages is drawn to scale. The serotypes of the host strains are indicated in parentheses. Amino acid sequence homologies are shown by shading with a heatmap.

**Fig. S6.** Procedures to determine the integration sites and genome sequences of Stx phages in O145:H28 strains with only draft genomes available. (A) Determination of integration sites by BLASTN search. The draft genomes of the O145:H28 strains (n=54) were searched by BLASTN using six query sequences: the *attL*-flanking and *attR*-flanking sequences from each of the prophage-integrated *wrbA* loci in strain 95-3192 (*wrbA*_L/R), the prophage-integrated *argW* locus in strain 122715 (*argW*_L/R), and the prophage-integrated *sbcB* locus in strain RM13514 (*sbcB*_L/R). Each query sequence was composed of the *att* sequence (7 bp for the prophage at *wrbA*, 24 bp for the prophage at *argW* and 12 bp for the prophage at *sbcB*) and the host chromosome and prophage sequences (60 bp each) that flanked the *att* sequence. Phage integration at each locus was considered positive when both *attL*– and *attR*-flanking sequences were detected (identity threshold: >95%). The integration sites of Stx phages in all but one genome were determined by this procedure. In strain IB14005, we detected the *argW*_L sequence but not the *argW*_R sequence. Therefore, the *argW* locus of this strain was defined as ‘Others’ and subjected to long PCR analysis. (B) Strategies for long PCR and the sequence determination of Stx phage genomes are shown. Stx prophage regions in each strain were divided into two or three segments and amplified by long PCR as indicated, and PCR products obtained from each strain were subjected to Illumina sequencing to determine the sequence of the entire prophage region. The complete sequences of two non-Stx phages integrated into *wrbA*, the sequences of the early regions of five non-Stx phages integrated into *sbcB*, and the full-length sequences of six degraded non-Stx phages integrated into *sbcB* were also determined by Type III, Type VIIa, and Type VIIb strategies, respectively.

**Fig. S7.** Optimization of the Stx2 production assay. (A) Lysis curves and Stx2 concentrations of seven O145:H28 strains (three strains carrying S-Stx2a phages and four strains carrying L-Stx2 phages). For each strain, the MMC-induced lysis curve (left) and the Stx2 concentrations in cell lysates obtained at each time point (right) are shown. Bacterial cells were inoculated into 40 ml of LB at an OD_600_ of 0.1 and grown to mid-log phage at 37 °C with shaking. MMC was added to the culture at final concentrations of 0.5, 1.0, 2.0, or 4.0 µg/ml. After the addition of MMC, the OD_600_ of each culture was measured every hour for 8 h, and 100 µl of the culture was collected at each time point to prepare cell lysates. The Stx2 concentration in each lysate was determined by sandwich ELISAs (n=1). In most cases, the maximum cell lysis and the highest Stx2 concentration were observed at 6 h. (B) Minimum inhibitory concentrations (MICs) of various *E. coli* strains against MMC were determined. The MMC susceptibilities of seven O145:H28 strains, five STEC strains belonging to other major STEC serotypes, and strains K-12 and ATCC25922 were determined according to the standard protocol outlined in Clinical and Laboratory Standards Institute (CLSI) guidelines (M100 Performance Standards for Antimicrobial Susceptibility Testing, 28th Edition, CLSI 2018). The MICs of all the tested strains were 2.0 µg/ml or 4.0 µg/ml.

**Table S1.** O145:H28 strains analyzed in this study.

**Table S2.** K-12 lysogens carrying Stx2a phages of O145:H28 strains.

**Table S3.** STEC strains carrying Stx phages that encode CI-4– or CI-5-type CI repressors.

**Table S4.** Primers used for long PCR amplification of prophage regions.

**Table S5.** Primers used to generate Stx phage deletion mutants.

## Reference

1. Scheutz F, Teel LD, Beutin L, Piérard D, Buvens G, Karch H, et al. Multicenter evaluation of a sequence-based protocol for subtyping Shiga toxins and standardizing Stx nomenclature. J Clin Microbiol. 2012;50:2951–2963.

2. Koutsoumanis K, Allende A, Alvarez-Ordóñez A, Bover-Cid S, Chemaly M, et al. Pathogenicity assessment of Shiga toxin-producing *Escherichia coli* (STEC) and the public health risk posed by contamination of food with STEC. EFSA j. 2020;18:5967

3. Ogura Y, Ooka T, Asadulghani, Terajima J, Nougayrède J-P, Kurokawa K, et al. Extensive genomic diversity and selective conservation of virulence-determinants in enterohemorrhagic *Escherichia coli* strains of O157 and non-O157 serotypes. Genome Biol. 2007;8:1.

4. Steyert SR, Sahl JW, Fraser CM, Teel LD, Scheutz F, Rasko DA. Comparative genomics and *stx* phage characterization of LEE-negative Shiga toxin-producing *Escherichia coli*. Front Cell Infect Microbiol. 2012;2:133.

5. Fuller CA, Pellino CA, Flagler MJ, Strasser JE, Weiss AA. Shiga toxin subtypes display dramatic differences in potency. Infect Immun. 2011;79:1329–1337.

6. Bielaszewska M, Mellmann A, Bletz S, Zhang W, Köck R, Kossow A, et al. Enterohemorrhagic *Escherichia coli* O26:H11/H-: a new virulent clone emerges in Europe. Clin Infect Dis. 2013;56:1373–1381.

7. Delannoy S, Mariani-Kurkdjian P, Webb HE, Bonacorsi S, Fach P. The Mobilome; A major contributor to *Escherichia coli stx2*-positive O26:H11 strains intra-serotype diversity. Front Microbiol. 2017;8:1625.

8. Yara DA, Greig DR, Gally DL, Dallman TJ, Jenkins C. Comparison of Shiga toxin-encoding bacteriophages in highly pathogenic strains of Shiga toxin-producing *Escherichia coli* O157:H7 in the UK. Microb Genom. 2020;16:16663.

9. Plunkett G, Rose DJ., Durfee TJ., Blattner FR. Sequence of Shiga toxin 2 Phage 933W from *Escherichia coli* O157:H7: Shiga toxin as a phage late-gene product. J Bacteriol. 1999;181:1767–1778.

10. Teel LD, Melton-Celsa AR, Schmitt CK, O’Brien AD. One of two copies of the gene for the activatable shiga toxin type 2d in *Escherichia coli* O91:H21 strain B2F1 is associated with an inducible bacteriophage. Infect Immun. 2002;70:4282–4291.

11. Muniesa M, Blanco JE, de Simon M, Serra-Moreno R, Blanch AR, Jofre J. Diversity of *stx2* converting bacteriophages induced from Shiga-toxin-producing *Escherichia coli* strains isolated from cattle. Microbiology. 2004;150: 2959–2971.

12. Strauch E, Schaudinn C, Beutin L. First-time isolation and characterization of a bacteriophage encoding the Shiga toxin 2c variant, which is globally spread in strains of *Escherichia coli* O157. Infect Immun. 2004;72:7030–7039.

13. Asadulghani M, Ogura Y, Ooka T, Itoh T, Sawaguchi A, Iguchi A, et al. The defective prophage pool of *Escherichia coli* O157: prophage-prophage interactions potentiate horizontal transfer of virulence determinants. PLoS Pathog. 2009;5:e1000408.

14. García-Aljaro C, Muniesa M, Jofre J, Blanch AR. Genotypic and phenotypic diversity among induced, stx2-carrying bacteriophages from environmental Escherichia coli strains. Appl Environ Microbiol. 2009;75:329–336.

15. Bonanno L, Petit M-A, Loukiadis E, Michel V, Auvray F. Heterogeneity in induction level, infection ability, and morphology of Shiga toxin-encoding phages (Stx phages) from dairy and human Shiga toxin-producing *Escherichia coli* O26:H11 isolates. Appl Environ Microbiol. 2016;82:2177–2186.

16. Pinto G, Sampaio M, Dias O, Almeida C, Azeredo J, Oliveira H. Insights into the genome architecture and evolution of Shiga toxin encoding bacteriophages of *Escherichia coli*. BMC Genomics. 2021;22:366.

17. Neely MN, Friedman DI. Arrangement and functional identification of genes in the regulatory region of lambdoid phage H-19B, a carrier of a Shiga-like toxin. Gene. 1998;223:105–113.

18. Makino K, Yokoyama K, Kubota Y, Yutsudo CH, Kimura S, Kurokawa K, et al. Complete nucleotide sequence of the prophage VT2-Sakai carrying the verotoxin 2 genes of the enterohemorrhagic *Escherichia coli* O157:H7 derived from the Sakai outbreak. Genes Genet Syst. 1999;74:227–239.

19. Strauch E, Hammerl JA, Konietzny A, Schneiker-Bekel S, Arnold W, Goesmann A, et al. Bacteriophage 2851 is a prototype phage for dissemination of the Shiga toxin variant gene 2c in Escherichia coli O157:H7. Infect Immun. 2008;76:5466–5477.

20. Fagerlund A, Aspholm M, Węgrzyn G, Lindbäck T. High diversity in the regulatory region of Shiga toxin encoding bacteriophages. BMC Genomics. 2022;23:230.

21. Yokoyama K, Makino K, Kubota Y, Watanabe M, Kimura S, Yutsudo CH, et al. Complete nucleotide sequence of the prophage VT1-Sakai carrying the Shiga toxin 1 genes of the enterohemorrhagic *Escherichia coli* O157:H7 strain derived from the Sakai outbreak. Gene. 2000;258:127–139.

22. Shi T, Friedman DI. The operator-early promoter regions of Shiga-toxin bearing phage H-19B. Mol Microbiol. 2001;41:585–599.

23. Calderwood SB, Mekalanos JJ. Iron regulation of Shiga-like toxin expression in *Escherichia coli* is mediated by the fur locus. J Bacteriol. 1987;169:4759–4764.

24. Wagner PL, Livny J, Neely MN, Acheson DWK, Friedman DI, Waldor MK. Bacteriophage control of Shiga toxin 1 production and release by *Escherichia coli*. Mol Microbiol. 2002;44:957–970.

25. Yano B, Taniguchi I, Gotoh Y, Hayashi T, Nakamura K. Dynamic changes in Shiga toxin (Stx) 1 transducing phage throughout the evolution of O26:H11 Stx-producing *Escherichia coli*. Sci Rep. 2023;13:4935.

26. Wagner Patrick L., Neely Melody N., Zhang Xiaoping, Acheson David W. K., Waldor Matthew K., Friedman David I. Role for a phage promoter in Shiga toxin 2 expression from a pathogenic *Escherichia coli* strain. J Bacteriol. 2001;183:2081–2085.

27. Tyler JS, Mills MJ, Friedman DI. The operator and early promoter region of the Shiga toxin type 2-encoding bacteriophage 933W and control of toxin expression. J Bacteriol. 2004;186:7670–7679.

28. Ogura Y, Mondal SI, Islam MR, Mako T, Arisawa K, Katsura K, et al. The Shiga toxin 2 production level in enterohemorrhagic *Escherichia coli* O157:H7 is correlated with the subtypes of toxin-encoding phage. Sci Rep. 2015;5:16663.

29. Nishida R, Nakamura K, Taniguchi I, Murase K, Ooka T, Ogura Y, et al. The global population structure and evolutionary history of the acquisition of major virulence factor-encoding genetic elements in Shiga toxin-producing *Escherichia coli* O121:H19. Microb Genom. 2021;7:e000716.

30. Miyata T, Taniguchi I, Nakamura K, Gotoh Y, Yoshimura D, Itoh T, et al. Alteration of a Shiga toxin-encoding phage associated with a change in toxin production level and disease severity in *Escherichia coli*. Microb Genom. 2023;9:e000935.

31. Nakamura K, Seto K, Lee K, Ooka T, Gotoh Y, Taniguchi I, et al. Global population structure, genomic diversity and carbohydrate fermentation characteristics of clonal complex 119 (CC119), an understudied Shiga toxin-producing *E. coli* (STEC) lineage including O165:H25 and O172:H25. Microb Genom. 2023;9:e000959.

32. Valilis E, Ramsey A, Sidiq S, DuPont HL. Non-O157 Shiga toxin-producing *Escherichia coli*-A poorly appreciated enteric pathogen: Systematic review. Int J Infect Dis. 2018;76:82– 87.

33. Nakamura K, Murase K, Sato MP, Toyoda A, Itoh T, Mainil JG, et al. Differential dynamics and impacts of prophages and plasmids on the pangenome and virulence factor repertoires of Shiga toxin-producing *Escherichia coli* O145:H28. Microb Genom. 2020;6:e000323.

34. Tyson S, Peterson C-L, Olson A, Tyler S, Knox N, Griffiths E, et al. Eleven high-quality reference genome sequences and 360 draft assemblies of Shiga toxin-producing *Escherichia coli* isolates from human, food, animal, and environmental sources in Canada. Microbiol Resour Announc. 2019;8:e00625–19.

35. Carter MQ, Pham A, Du W-X, He X. Differential induction of Shiga toxin in environmental *Escherichia coli* O145:H28 strains carrying the same genotype as the outbreak strains. Int J Food Microbiol. 2021;339:109029.

36. Nakamura K, Ogura Y, Gotoh Y, Hayashi T. Prophages integrating into prophages: A mechanism to accumulate type III secretion effector genes and duplicate Shiga toxin-encoding prophages in *Escherichia coli*. PLoS Pathog. 2021;17:e1009073.

37. Ondov BD, Treangen TJ, Melsted P, Mallonee AB, Bergman NH, Koren S, et al. Mash: fast genome and metagenome distance estimation using MinHash. Genome Biol. 2016;17:132.

38. Pacheco AR, Sperandio V. Shiga toxin in enterohemorrhagic *E.coli*: regulation and novel anti-virulence strategies. Front Cell Infect Microbiol. 2012;2:81.

39. Wagner PL, Acheson DW, Waldor MK. Human neutrophils and their products induce Shiga toxin production by enterohemorrhagic *Escherichia coli*. Infect Immun. 2001;69:1934– 1937.

40. Hall G, Kurosawa S, Stearns-Kurosawa DJ. Dextran sulfate sodium colitis facilitates colonization with Shiga toxin-producing *Escherichia coli*: a novel murine model for the study of Shiga toxicosis. Infect Immun. 2018;86:e00530–18.

41. Persson S, Olsen KEP, Ethelberg S, Scheutz F. Subtyping method for *Escherichia coli* Shiga toxin (verocytotoxin) 2 variants and correlations to clinical manifestations. J Clin Microbiol. 2007;45:2020–2024.

42. Dallman TJ, Ashton PM, Byrne L, Perry NT, Petrovska L, Ellis R, et al. Applying phylogenomics to understand the emergence of Shiga-toxin-producing *Escherichia coli* O157:H7 strains causing severe human disease in the UK. Microb Genom. 2015;1:e000029.

43. Cooper KK, Mandrell RE, Louie JW, Korlach J, Clark TA, Parker CT, et al. Comparative genomics of enterohemorrhagic *Escherichia coli* O145:H28 demonstrates a common evolutionary lineage with *Escherichia coli* O157:H7. BMC Genomics. 2014;15:17.

44. Patel PN, Lindsey RL, Garcia-Toledo L, Rowe LA, Batra D, Whitley SW, et al. High-quality whole-genome sequences for 77 Shiga toxin-producing *Escherichia coli* strains generated with PacBio sequencing. Genome Announc. 2018;6:e00391–18.

45. Wick RR, Judd LM, Gorrie CL, Holt KE. Unicycler: Resolving bacterial genome assemblies from short and long sequencing reads. PLoS Comput Biol. 2017;13:e1005595.

46. Croucher NJ, Page AJ, Connor TR, Delaney AJ, Keane JA, Bentley SD, et al. Rapid phylogenetic analysis of large samples of recombinant bacterial whole genome sequences using Gubbins. Nucleic Acids Res. 2015;43:e15.

47. Kurtz S, Phillippy A, Delcher AL, Smoot M, Shumway M, Antonescu C, et al. Versatile and open software for comparing large genomes. Genome Biol. 2004;5:R12.

48. Stamatakis A. RAxML-VI-HPC: maximum likelihood-based phylogenetic analyses with thousands of taxa and mixed models. Bioinformatics. 2006;22:2688–2690.

49. Tanizawa Y, Fujisawa T, Nakamura Y. DFAST: a flexible prokaryotic genome annotation pipeline for faster genome publication. Bioinformatics. 2018;34:1037–1039.

50. Ohtsubo Y, Ikeda-Ohtsubo W, Nagata Y, Tsuda M. GenomeMatcher: A graphical user interface for DNA sequence comparison. BMC Bioinformatics. 2008;9:1–9.

51. Nagano DS, Taniguchi I, Ono T, Nakamura K, Gotoh Y, Hayashi T. Systematic analysis of plasmids of the *Serratia marcescens* complex using 142 closed genomes. Microb Genom. 2023;9:e001135.

52. Kumar S, Stecher G, Li M, Knyaz C, Tamura K. MEGA X: Molecular evolutionary genetics analysis across computing platforms. Mol Biol Evol. 2018;35:1547–1549.

53. Islam MR, Ogura Y, Asadulghani, Ooka T, Murase K, Gotoh Y, et al. A sensitive and simple plaque formation method for the Stx2 phage of *Escherichia coli* O157:H7, which does not form plaques in the standard plating procedure. Plasmid. 2012;67:227–235.

54. Ooka T, Terajima J, Kusumoto M, Iguchi A, Kurokawa K, Ogura Y, et al. Development of a multiplex PCR-based rapid typing method for enterohemorrhagic *Escherichia coli* O157 strains. J Clin Microbiol. 2009;47:2888–2894.

55. Datsenko KA, Wanner BL. One-step inactivation of chromosomal genes in *Escherichia coli* K-12 using PCR products. Proc Natl Acad Sci USA. 2000;97:6640–6645.

56. Nakamura K, Tokuda C, Arimitsu H, Etoh Y, Hamasaki M, Deguchi Y, et al. Development of a homogeneous time-resolved FRET (HTRF) assay for the quantification of Shiga toxin 2 produced by E. coli. PeerJ. 2021;9:e11871.

57. R Core Team. R: A language and environment for statistical computing. manual, Vienna, Austria. 2020.

